# Taxonomic revision of Leopold and Rudolf Blaschka Glass Models of Invertebrates 1888 Catalogue, with correction of authorities

**DOI:** 10.1101/408260

**Authors:** Eric Callaghan, Bernhard Egger, Hazel Doyle, Emmanuel G. Reynaud

**Author notes:** Correspondence should be addressed to: Conway Institute School of Biomolecular and Biomedical Science University College Dublin, Belfield, Dublin 4, Ireland. ORCID: 0000-0003-1502-661X.

## Abstract

The glass models of invertebrates crafted by Leopold and Rudolf Blaschka were made between 1863 and 1889. Production ceased when the glassmakers turned their attention to what is now known as the Ware Collection of Blaschka Glass Models of Plants, created for the Harvard Museum of Natural History. Nearly 130 years have passed since their last published catalogue of species in 1888 and the nomenclature they applied is now a confusing mix that includes many junior synonyms and unavailable names. This is an issue for many museums and universities which owns Blaschka models as uncertain identifications may compromised interpretation of this rediscovered legacy. Nowadays, many museums and universities hold collections of those glass invertebrates but misspelled, mislabelled or simply apply the century old taxonomy of 1888. Here, we provide a valuable resource for curators and enthusiasts alike. We studied and updated the final catalogue 1888 from the Blaschkas’ Dresden-based workshop. We first focused on major taxonomical changes from taxa to species, as well as on an analysis of the acknowledged authorities. We found that only 35.3% of the models retain their original names, while 3.7% lack any known synonym and their identity remains open to interpretation. Finally, two of the authorities listed in the catalogue, Ernst Haeckel and Philip Henry Gosse, were incorrectly acknowledged for an extensive range of models. This study is the first of its kind on the Blaschka collection, and it will help in the identification and naming of Blaschka models worldwide.

## INTRODUCTION

During the 18th century, the Swedish botanist Carl von Linné (Carolus Linnaeus) established a “two-term naming system”, also known as the binomial nomenclature to name every specie. This system is now governed by international codes of rules such as the *International Code of Zoological Nomenclature* (ICZN). This system encompasses terrestrial as well as marine species that became the focus of many expeditions in the 19th century. The reports on these explorations, from François Auguste Péron’s jellyfish drawings (Péron, 1816) to Ernst Haeckel’s radiolarian engravings (Haeckel, 1887), alongside the massive 35 volumes from the HMS *Challenger* expedition reports (1872–1876), opened a new world to the masses, enabling them to see these creatures both in books and in prints, but also in museums and exhibitions. This museum movement, designed to educate the people and to display what the world has to offer, could easily be advanced by the display of skeletons and exotic stuffed animals, but the marine world, other than fishes and dolphins, remained difficult to display.

One workshop, based in the German town of Dresden, found a solution to the challenge of displaying the newly described marine invertebrates, which usually faded, or shrank in preservatives. The glassblowers Leopold and Rudolf Blaschka, father and son, used their knowledge of glass and its translucent qualities, as well as pigments to create artificial jellyfishes and other soft-bodied invertebrates that could be exhibited easily (Reiling; 1998 and 2000). However, they relied on books, lithographs, and sometimes live creatures kept in tanks to produce their models (Dohrn A.; 1877). Many different books and monographs were used as source illustrations such as Philip Henry Gosse’s *Actinologia Britannica: A History of the British Sea-Anemones and Corals* (Gosse, 1860), Haeckel’s *Das System der Medusen* (Haeckel, 1879) or Jean Baptiste Vérany’s *Céphalopodes de la Méditerranée* (Vérany, 1851). The Blaschkas created more than 700 models that were sold worldwide through their own workshop and also through three dealers: Robert Damon (United Kingdom and Ireland), Václav Frič (Austria and Hungary), and Henry Augustus Ward (North America). These models are quality representations, and they are often referred to as masterpieces in which their art matches their true biological nature (Sheets-Pyenson, 1988; Dyer, 2008).

Since the production of these magnificent models ended in 1889, a wealth of marine biological data has accumulated, and there have been many taxonomic changes. In addition, challenges to established ideas and concepts have led to the extensive reorganization of the Tree of Life (e.g., the Archean Kingdom). However, the name “glass models of invertebrates,” which has been consistently applied to the Blaschkas’ creations, has never been challenged, presumably because these models were extremely accurate, and little has been published about their taxonomy. Although some work has been done on the origin of their designs and their sources of inspiration, it is often very general and incomplete.

We decided to investigate the taxonomy of the Blaschkas’ 700-plus glass models of invertebrates listed in the two English catalogues (1878;1888) published by Ward’s Natural Science Establishment from archives such as those housed in the Rakow Research Library of The Corning Museum of Glass (which contains the archives the Blaschkas’ workshop), as well as the large, digitized holdings of the online Biodiversity Heritage Library (BHL) and World Register of Marine Species (WoRMS). We also analyzed the authority for each species and the taxonomic value of the original species’ name versus the currently established one. We thus established a new version of the Blaschkas’ catalogue with the correct modern taxonomy and authority for each species, along with a unique set of “Blaschka species” that exist only as glass models (the species they described are no longer considered valid). Finally, we uncovered a bias toward using the work of the British naturalist Philip Henry Gosse and Ernst Haeckel as recognized authorities.

## MATERIAL AND METHODS

### Archival Material

The two original catalogues that describe the glass invertebrate models sold by the Blaschkas’ workshop in Dresden were obtained from the following sources: Ward’s Natural Science Establishment catalogue (1878): Reese Library of the University of California. (online access: https://babel.hathitrust.org/); Ward’s Natural Science Establishment catalogue (1888): River Campus Libraries, University of Rochester, Rochester, New York, Henry Augustus Ward Papers (1840–1933), reference A.W23.

### Analysis of Data

Because of the extent of the species and phyla covered by the Leopold and Rudolf Blaschka, we had to work, for the most part, on well-established and curated online databases to ascertain the validity of the models. All the model specie names were checked, and the taxonomy—from phylum to species—was updated as much as possible.

The principal databases consulted were: World Register of Marine Species (WoRMS), www.marinespecies.org; Marine Species Identification Portal, species-identification.org; and the Catalogue of Life, www.catalogueoflife.org.

We also used the Biodiversity Heritage Library, www.biodiversitylibrary.org. This is a source of scanned original books that also contains chromolithographies and help us to compare with Blashcka drawings and the final modeland confirm or infirm the binomial nomenclature.

We always rely on the provided information from those databases. Depending on the final established taxonomy, we used the following provided taxonomical naming: “*nomen dubium*” (Latin, “doubtful name,” indicating that the taxonomic validity is uncertain or disputed by various experts), “*nomen nudum*” (Latin, “naked name,” indicating a name that has been published without an adequate description), and “*species inquirenda*” (Latin, “species of doubtful identity, requiring further investigation”). In cases where no matching entry could be found in any of these databases, an online search was conducted to cross-reference other sources, which often clarified the identification or suggested a possible alternative. For several models, no current identification could be found, despite our best efforts. These models are then designated as “ND” (No Data) in the updated version of the catalogue.

## RESULTS

### General Catalogue Analysis

There are two catalogues of the models produced by the Blaschka workshop, which were made available to us from different archives. In 1878, Henry Augustus Ward published a catalogue specifically for the Blaschkas’ invertebrate models, probably based on a German edition of the glassmakers’ workshop catalogue, in which the models are listed according to the generally accepted invertebrate taxonomy (Ward, 1878). Each model was assigned a unique catalogue number (1 to 630) that would become a standard reference. For ease of reference the models in this article, are ordered according to these numbers as these are consistent and reliable identifiers. Because this catalogue was produced in the United States of America, the models are priced in dollars and cents.

In 1888, Ward printed a revised catalogue to include the models that had been added by the Blaschkas since the publication of his 1878 catalogue, but new additions (631 to 704 + 6 non-numbered models) to this new catalogue were rather listed at the end. This means that many phyla have models appearing in two sections in the numbered list; once in Models 1-630 and again amongst 631-704. The first 630 models show a traditional textbook systematic arrangement for numbers 1-630 as per Ward 1878. Numbers 630-713 are models subsequently added to the repertoire in later catalogues and show a similar systematic ordering although with sponges added last.

The Blaschkas were glassblowers, not taxonomists, and they had to rely on the limited taxonomic literature available at the time and especially chromolithographies within them that allows them to produce coloured models. The most well-known example is the anemones based lithograpies illustrated P.-H. Gosse book (e.g. *Caryophilla Smithii*). Henry Ward was a geologist. At that time, it was customary to assign a specific status to organisms based on minor differences that would today be regarded as a subspecies at best, and therefore some of the models in the catalogues represent “species” that are no longer considered valid. In addition, it is possible that some of the species were incorrectly identified in the first place.

None of the two catalogues follow the established taxonomic conventions, in that the generic and specific names are not italicized. Also, species names were capitalized when they refer to persons which was common practice in the literature of the time (e.g., Model no. 30, *Actinoloba Paumotensis*, and Model no. 43, *Bunodes Ballii*).

There are many spelling errors throughout the catalogues. . These may have been the fault of the printers, who would not have been experts in the field (e.g., model no. 20 is listed as *Actinaria* rather than *Actiniaria*). These mistakes may indicate that neither Ward nor the Blaschkas corrected their manuscripts before they were printed.

### Analyzing Ward’s 1888 Catalogue

We used Henry Ward’s 1888 catalogue as the last available catalogue to establish a reference of the complete Blaschka marine invertebrate collection. Seven hundred four models are perfectly numbered, and there are six models at the end of the catalogue that have no numbers, establishing 713 as the complete set of models offered to customers. However, the distribution is highly variable across phyla, classes, and orders (Table 1). Moreover, there are minor discrepancies in the catalogue. Model 141 (*Cladonema radiatum*) is listed twice (development and adult), but there is only one associated number because the model of the adult is not numbered separately. The same situation applies for Model 191 (*Tubularia indivisa*). Also, in addition to the model numbered 219 (*Rhizophysa Eysenhardti*, there is a model 219a (*Rhizophysa heliantha*, now *Anthorhybia rosacea*).

**TABLE 1:**
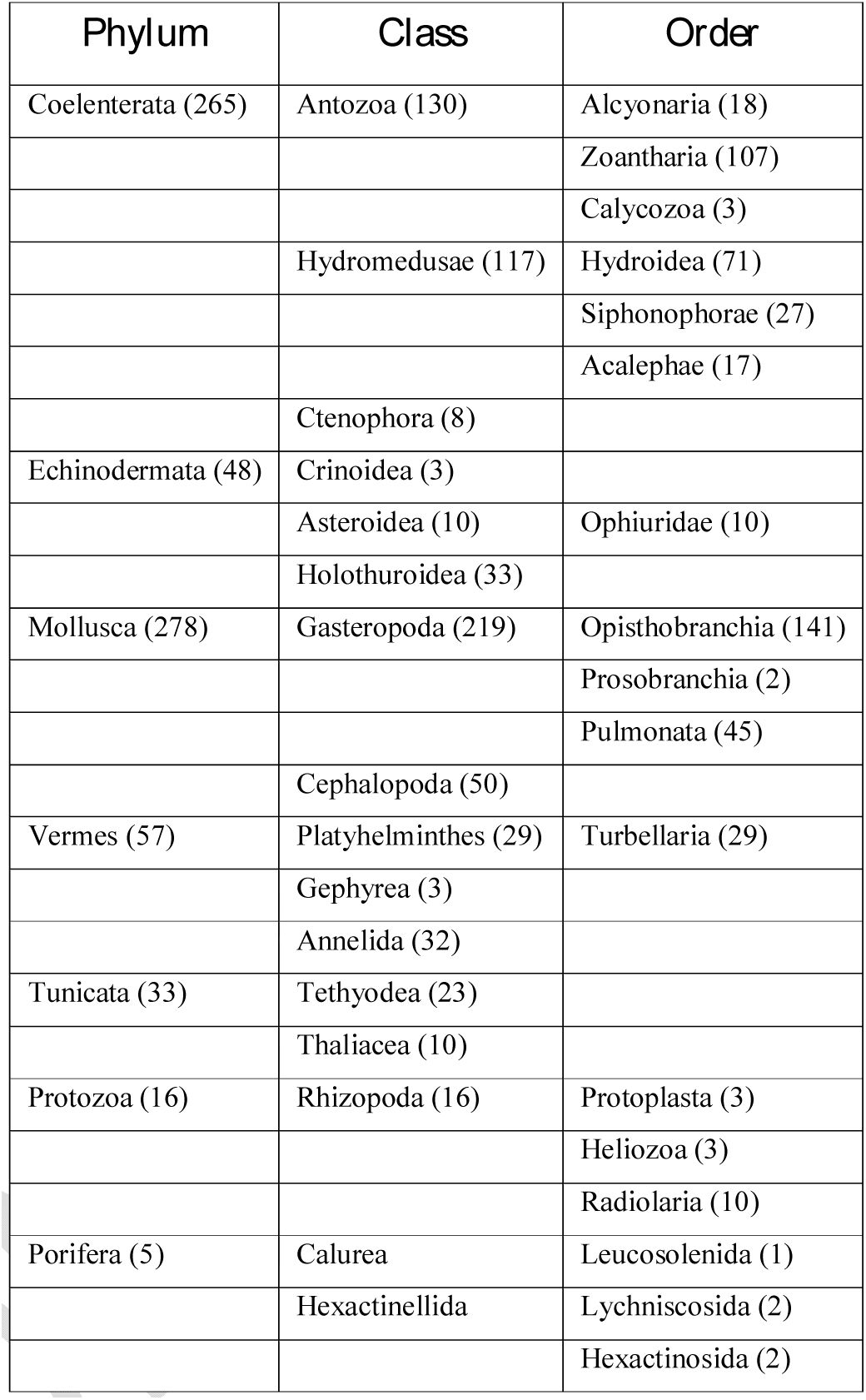
Taxonomic Distribution of Invertebrate Models in Henry Ward’s 1888 Catalogue

Of the 713 models, 19 (2.6%) are of varieties no longer considered valid, although three of these are now regarded as full species in their own right; 10 (1.4%) represent developmental stages of species (note that models 252 and 669 have no adult forms listed in the catalogue); 12 (1.7%) illustrate the anatomy of species, three of which are not otherwise included in the catalogue; and four (0.6%) represent male and female specimens of two species. Therefore, the 713 models represent 694 species *as recognized at that time.*

### General Changes in Taxonomy (from the 1888 Ward Catalogue)

At the phylum level, three phyla are still valid (Echinodermata, Mollusca, and Porifera) and two phyla (Coelenterata and Vermes) are obsolete, while Tunicata is now a subphylum of Chordata. The Protozoa, introduced in 1818 as a taxonomic class, has been and remains a problematic area of taxonomy but is currently considered a subkingdom in the kingdom Protista. Coelenterata now encompasses the current phyla Ctenophora (comb jellies) and Cnidaria. Platyhelminthes and Annelida are now two phyla that cover the obsolete Vermes phylum. (In the catalogues, the term “Phylum” does not appear; instead, the now obsolete “Type” is found.)

At the Class level, eight classes are still correct (Anthozoa, Crinoidea, Asteroidea, Holothuroidea, Gastropoda (originally Gasteropoda), Cephalopoda, Thaliacea, and Turbellaria), and one is obsolete (Gephyrea). However, because of the reorganization of phyla and subphyla, many classes are now assigned to various phyla and subphyla (e.g., Anthozoa is now a class of the phylum Cnidaria) (Table 2). Three classes used name that can be commonly found with different spellings: Hydromedusae (Hydroidomedusae, now accepted as Hydroidolina), Gasteropoda (Gastropoda), and Tethyodea (Tethioidea). This could be based on the original book used for the specie name or eventually some printing errors or transcription.

**TABLE 2:** Corrected Taxonomic Distribution at the Class and Order Levels of Marine Invertebrate Models in the 1888 Ward Catalogue

| *Class* | *Current Status/Rank* | *Comments* |
| --- | --- | --- |
| Anthozoa | Class | Class in Phylum Cnidaria |
| Hydromedusae<br>(Hydroidomedusae) | Class (Hydroidolina) | Subclass of Hydrozoa, phylum Cnidaria |
| Crinoidea | Class | Class in Subphylum Crinozoa, phylum Echinodermata |
| Asteroidea | Class | Class in Subphylum Asterozoa, phylum Echinodermata |
| Holothuroidea | Class | Class in Subphylum Echinozoa, phylum Echinodermata |
| Gasteropoda | Class (Gastropoda) | Class in Phylum Mollusca |
| Cephalopoda | Class | Class in Phylum Mollusca |
| Gephyrea | Obsolete |  |
| Tethyodea<br>(Tethioidea) | Division | Division of Subphylum Tunicata |
| Thaliacea | Class | Class of Subphylum Tunicata |
| Turbellaria | Class | Class in Phylum Platyhelminthes |
| Alcyonaria | Subclass (Octocorallia) | Subclass of Anthozoa |
| Zoantharia | Order | Order of Subclass Hexacorallia, class Anthozoa |
| Calycozoa | Obsolete |  |
| Hydroidea | Obsolete |  |
| Siphonophorae | Order | Order of Class Hydrozoa |
| Acalephae | Obsolete |  |
| Ophiuridae | Family | Family of Order Ophiurida |
| Opisthobranchia | Infraclass | Infraclass of Class Gastropoda |
| Prosobranchia | Subclass | Infraclass of Class Gastropoda (Prosobranchia is no longer accepted as a valid subclass see Ponder & Lindberg, 1997) |
| Pulmonata | Infraclass | Infraclass of Subclass Heterobranchia |
| Turbellaria | Class | Class in Phylum Platyhelminthes |

At the Order level, there have been extensive changes, as noted in Table 2. Three orders are now obsolete (Calycozoa, Hydroidea, and Acalephae), while many orders are now regarded as classes, infraclasses, subclasses, or families. Only two orders remain valid today (Zoantharia and Siphonophorae).

Concerning the Species taxonomic classification of the Blaschka marine invertebrate models, 240 (33.7%) are unchanged, 400 (56.1%) have changed (this includes the variations that are no longer recognized), and 40 (5.6%) have been only tentatively identified. For 25 (3.5%), no data can be located (this includes one model that bears the name of a plant species). Finally, four (0.56%) are described as “*nomen dubium*,” two (0.28%) are termed “*nomen nudum*,” and two (0.28%) are regarded as “*species inquirenda*.”

Interestingly, 60 models (8.4% of the catalogue) are of species that had been described within the preceding 30 years (i.e., since 1858), and 17 of those (2.4% of the catalogue) had been described within the preceding 20 years (i.e., since 1868).

### Authority

In taxonomic nomenclature, it is common practice to identify a species using the established binomial name, followed by the “authority.” It is a way of identifying the person who first published the name, and it is a very important component of the species’ nomenclature. We identified 136 naming authorities, but 22 of these accounted for 64 percent of the names. They include such well-recognized naturalists as Carl von Linné and Jean-Baptiste Lamarck, but also some authors who are regarded as experts in specific branches of invertebrate studies: Louis Agassiz and Edward Forbes (Cnidaria), Jacques Philippe Raymond Draparnaud (Gastropoda), and Otto Friedrich Müller (Actiniaria).

Philip Henry Gosse, the English naturalist and popular nature writer, is the principal naming authority quoted, with 59 species in the catalogue attributed to him. However, the identification of 50 of these species has been revised. Twelve were reassigned to species already described by Gosse, and 38 were reclassified as species previously identified by other authorities. Only nine were retained as genuinely new species described by Gosse.

Another frequently quoted authority is Ernst Haeckel. Twenty-one species are attributed to Haeckel in the catalogue, 13 of which have been reidentified (four as species previously described by Haeckel, and nine as species previously identified by other authorities). The remaining eight are unchanged as genuinely new species described by Haeckel.

## DISCUSSION

The Blaschka workshop, based in Dresden, developed a unique series of marine invertebrate models between 1863 and 1890, using as reference zoological illustrations such as those contained in Gosse’s *Actinologia Britannica* or Ludwig Schmarda’s *Neue wirbellose Thiere* (1859–1861). Although the current focus by many museums and universities on Blaschka models is welcome for highlighting invertebrate biology, interpretation of this rediscovered legacy is compromised by uncertain identifications. With the passing of time and new expeditions and discoveries, the extent of knowledge of the biological world increased, as did the complexity of the Tree of Life and the taxonomic keys required to identify every single species.

Because the Blaschkas’ models have always been regarded as accurate representations of invertebrates, we investigated the taxonomy of their entire production (713 models) to understand the zoological accuracy of their workshop and the development in understanding of those species over the last 127 years (1888–2015). We established the modern taxonomy of as many models as possible to provide every Blaschka collection curator with a reference table (Table 3) to properly label models with accurate taxonomic identification. But this table will not be the final one because we still have a series of models for which only limited information can be located. Forty models (5.6%) have been only tentatively identified (Table 4), no data can be located for 25 others (3.5%) (Table 4), four (0.56%) are described as “*nomen dubium*,” two (0.28%) are termed “*nomen nudum*,” and two (0.28%) are “*species inquirenda*” (Table 5). These will require further research.

**TABLE 3:** Complete Corrected Taxonomic Naming and Authority of Marine Invertebrate Models in Ward’s 1888 Catalogue

**[?]:** Identification that may require confirmation pending further investigation**; *Species inquirenda***: species the taxonomic validity of which is uncertain or disputed by different experts; ***Nomen nudum***: has not been published with an adequate description; ***Nomen dubium*:** a name of uncertain taxonomic significance; **Morpha**: Taxon below the species level, that has no taxonomic status; it consists of specimens in a population that differ in morphology from other group(s) of specimens in the same population; **MSIP** Marine Species Identification Portal; **WoRMS** World Register of Marine Species; **ND** No Data

**TABLE 4:**
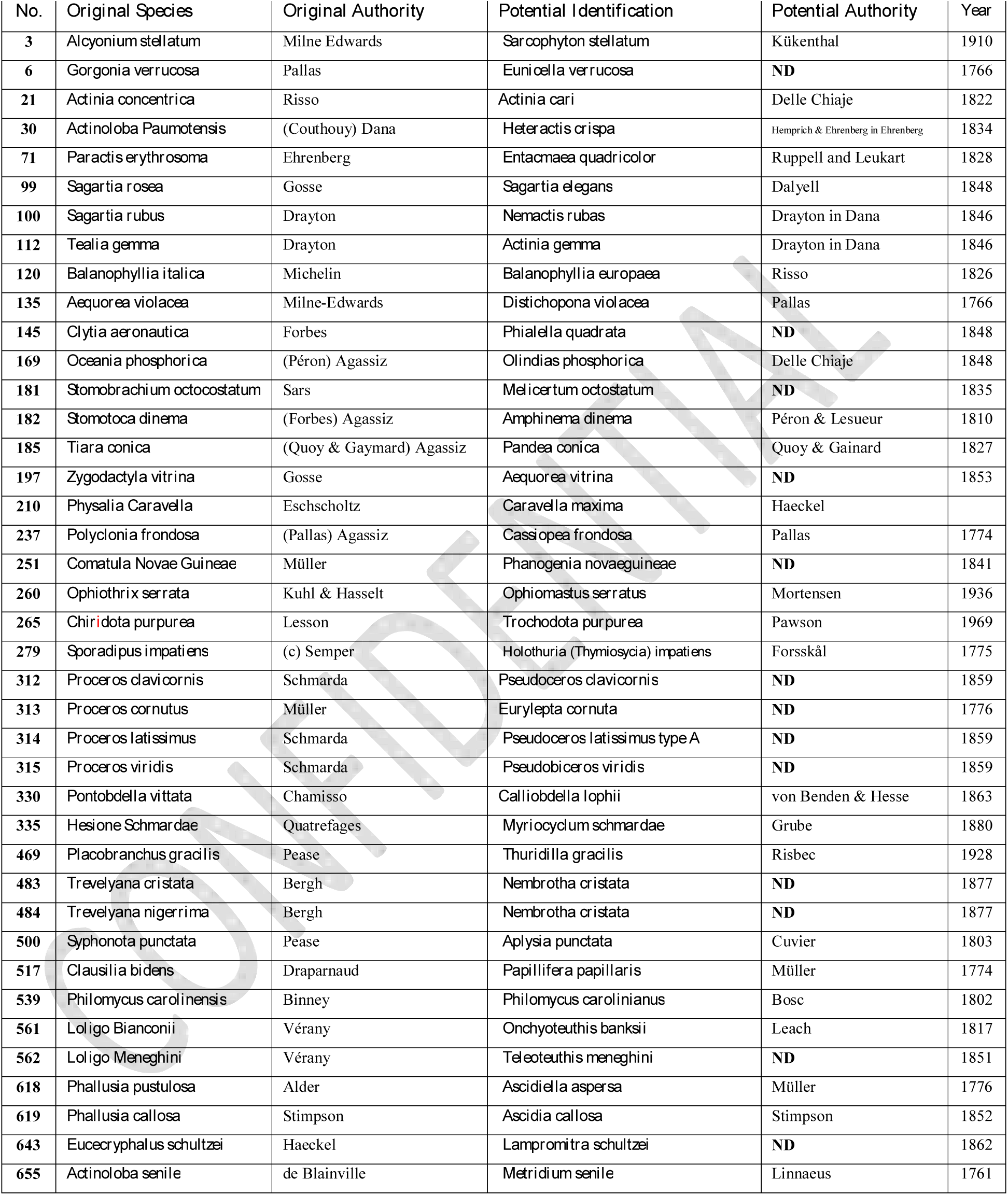
Species with uncertain or tentative identifications

**TABLE 5:** Species with No Identification Information

| **No.** | **Original Species Name** | **Authority** |
| --- | --- | --- |
| 12 | *Renilla violacea* | Quoy & Gaimard |
| 15 | *Sympodium purpurascens* | Ehrenberg |
| 60 | *Edwardsia vestita* | Forbes |
| 70 | *Paractis adhaerens* | Ehrenberg |
| 72 | *Paractis olivacea* | Ehrenberg |
| 87 | *Saccanthus purpurascens* | Milne Edwards |
| 148 | *Cunina campanulata* | Eschscholtz |
| 160 | *Liriope appendiculata* | Forbes |
| 168 | *Obelia sphaerulina* | Péron |
| 175 | *Polyxenia Alderii* | Forbes |
| 176 | *Rhegmatores (Aequorea) Forbesianus* | Gosse |
| 190 | *Trachynema ciliatum* | Gegenbaur |
| 194 | *Turris neglecta* | Forbes |
| 196 | *Zygodactyla crassa* | Agassiz |
| 198 | *Abyla pentagona* | Eschscholtz |
| 199 | *Agalma rigidum* | Haeckel |
| 207 | *Halistemma punctatum* | Kolliker |
| 209 | *Hippopodius gleba* | Leuckart |
| 211 | *Physalia pelagica* | Eschscholtz |
| 233 | *Holigoclados lunulatus* | Pennant |
| 368 | *Aeolis militaris* | Alder & Hancock |
| 392 | *Cratena longibursa* | Bergh |
| 442 | *Facellina Drummondii* | Thompson |
| 697 | *Paludina achatina* | Sowb |
|  | *Actinia chioocca* | Cocks |

It is interesting to note that, of the 630 models presented in the 1878 Ward catalogue and the 713 in Ward’s 1888 edition, we can identify only 694 species. Because of the invalidation of 25 variations of some species and the paucity of firm data on other species referred to in the preceding paragraph, we could finally retrieve only 621 valid and fully identified species, with 400 (64%) being unchanged since the catalogue of 1888. The Blaschkas’ workshop produced a series of models of developmental stages for some species (for example, Model no. 225, *Aurelia aurita*) and dissections illustrating the internal anatomy of others (for example, Model no. 275, *Holothuria tubulosa*). In each case, these models are listed in the catalogues under separate numbers, accounting for some of the apparent discrepancies between the number of models listed and the number of species they represent. There are more individual models than the catalogue numbering system implies because, in the species where the developmental stages have been produced, these are presented as a series of models grouped together in one display and listed under a single number.

Environmental conditions can exert a significant influence on the physical appearance of some species. In the past, it was common practice to describe and name animals and plants exhibiting these effects as distinct varieties within a species—a practice that is no longer considered valid. For example, model nos. 122, *Caryophyllia Smithii var. clara*, and 123, *var. castanea*, are no longer separated, but are listed as *Caryophyllia Smithii*.

One particularly interesting part of our findings is related to the naming authorities cited. In taxonomy, a species name is always linked to the name of the person who originally named it and the year when this occurred. Philip Henry Gosse had always been an important influence on the Blaschkas, father and son, as a well-established marine invertebrate expert, even though he was not a zoologist but rather a naturalist and popularizer of natural science. We have noted that the Blaschkas wrongly attributed many species (38 out of 59) to him. We have identified a similar situation with regard to another great influence on the workshop: Ernst Haeckel. We looked in detail at *Actinologia Britannica*, one of the major books known to have been used by the two glassworkers, and we found that the identification of the authority is often confusing for non-experts and may have been the source of the mistaken identities. In some instances, the Blaschkas listed Gosse himself as the naming authority, but Gosse did not list the actual naming authorities in his illustrations. Wherever a species can be clearly identified, we have retrieved the correct authority (Table 3).

Our work represents an important step toward establishing a complete descriptive database of the Blaschkas’ glass invertebrate models, enabling us to identify models and their names in accordance with both the original documents and current taxonomic knowledge. We have already performed several curations of European collections and corrected identification errors that were usually related to the loss of original labels or the mixing of those labels during curation, repair, or display. We expect that our Table 3 will be updated because more taxonomists will be able to access the relevant taxonomic information to confirm or correct the identification of the models, and to allow for the taxonomic identification of models for which we have no data (Table 5).

We will continue to use the information gathered during our research to link every model to the original documentation and lithograph used, alongside the drawings held at the Rakow Research Library of The Corning Museum of Glass. We believe that, although the Blaschkas’ invertebrate models are often described as unique art pieces, they were originally zoological specimens that need to be curated taxonomically and clearly identified and labeled, even if the species are no longer recognized. We hope that our work will help the Blaschka-related community to curate their collections in a taxonomically correct manner.

## ACKNOWLEDGEMENTS

We wish to thank the following for their help with this project: Nigel Monaghan and Paolo Viscardi at National Museum Ireland – Natural History.

## SECOND SUMMARY

Les modèles en verre d’invertébrés de Léopold et Rudolf Blaschka ont été fabriqués entre 1863 et 1889. Leur production a cessé lorsque ses deux maitres verriers ont tourné leur attention sur ce qu’on appelle désormais la collection Ware des modèles de plantes en verre, créés pour le Musée d’histoire naturelle de Harvard. Près de 130 ans se sont écoulés depuis leur dernier catalogue d’espèces d’invertébrés publié en 1888, et de nombreux événements se sont produits dans le monde de la biologie et de la taxonomie des invertébrés. De nos jours, de nombreux musées et universités possèdent des collections d’invertébrés Blaschka en verre, mais souvent mal orthographiés, mal étiquetés ou appliquant simplement la taxonomie centenaire de 1888. Nous fournissons ici une ressource précieuse pour les conservateurs et les passionnés. Nous avons étudié et mis à jour le catalogue final de 1888 de l’atelier de Blaschkas à Dresde. Nous nous sommes d’abord concentrés sur les changements taxonomiques majeurs des taxons aux espèces, ainsi que sur une analyse des autorités reconnues. Nous avons constaté que seulement 35,3% des modèles ont conservé leurs noms d’origine, tandis que 3,7% semblent avoir disparu. Enfin, deux des autorités énumérées dans le catalogue, Ernst Haeckel et Philip Henry Gosse, ont été mal attribués pour une vaste gamme de modèles. Cette étude est la première de son genre sur la collection Blaschka, et elle aidera à identifier et nommer les modèles Blaschka dans le monde entier.

## Weblinks

A. http://www.animalbase.unigoettingen.de/zooweb/servlet/AnimalBase/home/speciestaxon?id=15639

